# monaLisa: an R/Bioconductor package for identifying regulatory motifs

**DOI:** 10.1101/2021.11.30.470570

**Authors:** Dania Machlab, Lukas Burger, Charlotte Soneson, Filippo M. Rijli, Dirk Schübeler, Michael B. Stadler

## Abstract

Proteins binding to specific nucleotide sequences, such as transcription factors, play key roles in the regulation of gene expression. Their binding can be indirectly observed via associated changes in transcription, chromatin accessibility, DNA methylation and histone modifications. Identifying candidate factors that are responsible for these observed experimental changes is critical to understand the underlying biological processes. Here we present *monaLisa*, an R/Bioconductor package that implements approaches to identify relevant transcription factors from experimental data. The package can be easily integrated with other Bioconductor packages and enables seamless motif analyses without any software dependencies outside of R.

**Availability:** *monaLisa* is implemented in R and available on Bioconductor at https://bioconductor.org/packages/monaLisa with the development version hosted on GitHub at https://github.com/fmicompbio/monaLisa.

## Introduction

Binding proteins that interact with specific nucleotide sequences, such as transcription factors (TFs), play key roles in the regulation of cellular functions and organismal development (Spitz & Furlong, 2012). Identifying candidate proteins that could play regulatory roles in development or act as drivers for an observed biological response is thus a crucial step in the interpretation of genomics data, such as absolute values or changes of DNA methylation, chromatin modifications, accessibility or transcription. There are various existing tools and methods for regulatory protein identification via their binding motifs. Most of these are command line tools or web servers that cannot be easily integrated with other Bioconductor (Huber *et al.*, 2015) packages for a seamless analysis in R, or they require the installation of additional software outside of R. Conceptually, many of these methods can be roughly divided into two types: enrichment-based methods that compare motif occurrences between sets of sequences, and model-based methods that estimate motif importance from their ability to explain experimental observations. Here, we present *monaLisa*, short for “motif analysis with Lisa”, an R/Bioconductor package that implements both of these approaches and enables seamless motif identification analyses in R.

## Usage and examples

Enrichment-based tools like *HOMER* (Heinz *et al.*, 2010) and *MEME* (Bailey *et al.*, 2015) identify novel or known motifs enriched in a given set of sequences compared to a suitable background. In *monaLisa*, this is done by first binning sequences, for example gene promoters or enhancers, according to their associated values. In the example here we use changes of DNA methylation between mouse embryonic stem cells and derived neuronal progenitors (Stadler *et al.*, 2011; Burger *et al.*, 2013)(Fig. 1A). A collection of known motifs, for example from the *JASPAR2020* package (Fornes *et al.*, 2020), are then evaluated for enrichment in each bin compared to the background, using *HOMER*’s normalization method to adjust for differences in sequence composition. This can also be diagnosed using available visualization functions (Suppl. Fig. 1A and B). Several ways to define background sequences are available, and the results can be visualized as a heatmap (Fig. 1B), that shows the enrichment of each motif in each bin, compared to all other bins. Additional confidence can often be gained by focusing on motifs for which the enrichment scales with the numerical value under consideration. *monaLisa* also offers to search for enriched k-mers (oligonucleotides of length k), which is particularly useful to complement the motif enrichment analysis and identify potential gaps in the database of known motifs (Suppl. Fig. 1C).

**Figure 1:**
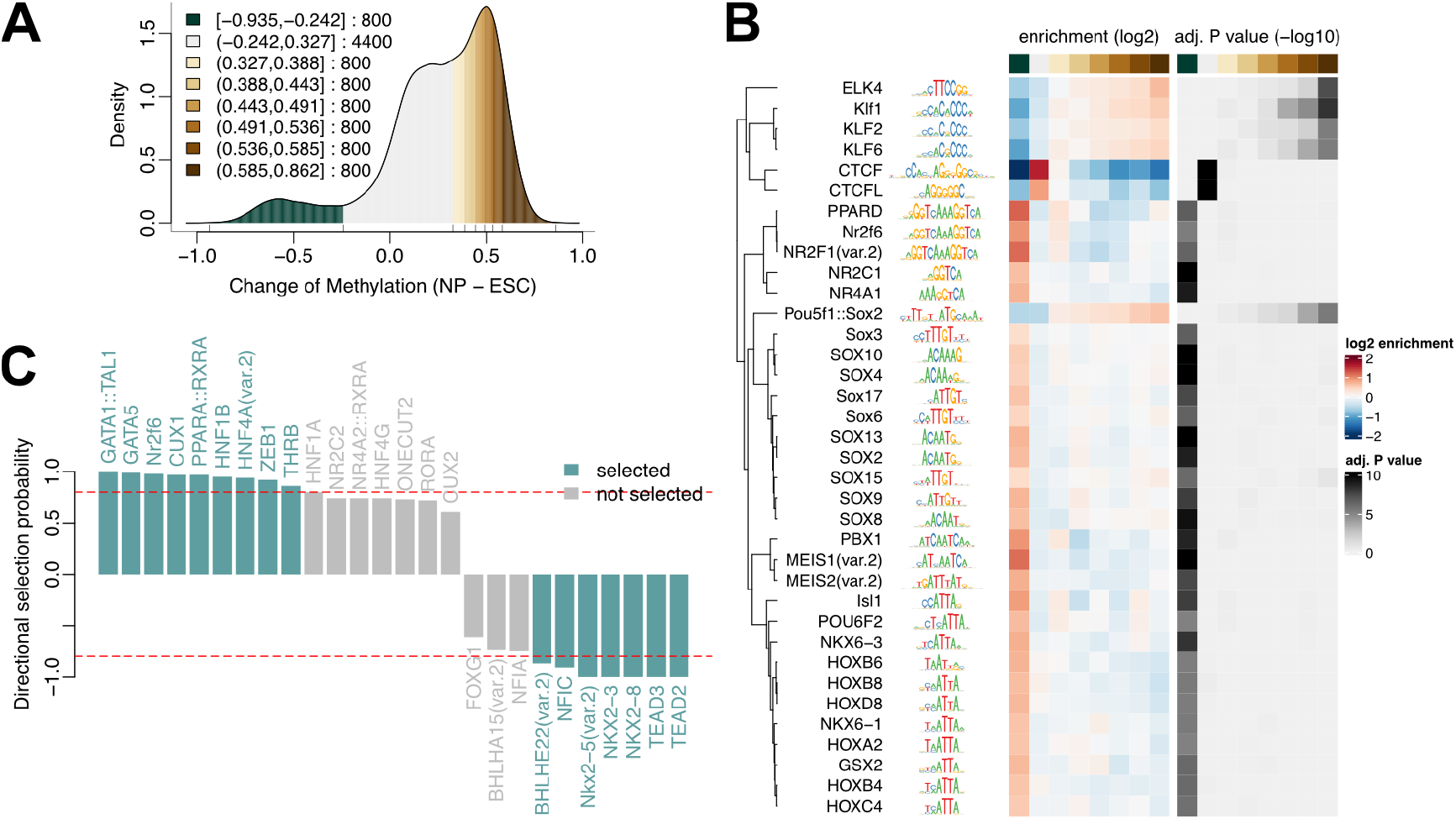
**A,B** Analysis of methylation changes between neuronal progenitors (NP) and embryonic stem cells (ESC): Binned density of methylation levels (A), with bin boundaries and sizes given in the legend, and enrichment and significance heatmaps (B) of motifs (rows) across bins (columns). **C** Analysis of accessibility changes between liver and lung: Directional selection probabilities for motifs identified using stability selection.

In the bin-based approach, motifs are analyzed independently of each other. In contrast, methods such as *REDUCE* (Roven & Bussemaker, 2003) or *ISMARA* (Balwierz *et al.*, 2014) use linear regression approaches to identify regulatory motifs that are most likely to explain the observed numerical responses. A similar model-based approach is also available in *monaLisa*, but uses a different regression framework: randomized lasso stability selection, introduced by Meinshausen & Bühlmann, 2010, with the improved error bounds proposed by Shah & Samworth, 2013. Regression is performed on several random subsets of the data to calculate motif selection probabilities. This type of regularization has advantages in selecting variables consistently, demonstrating better error control and not depending strongly on the initial regularization chosen (Meinshausen & Bühlmann, 2010). For illustration, we have used *monaLisa*’s regression with stability selection to identify transcription factor motifs that could explain the observed changes of accessibility between mouse liver and lung (data from The ENCODE Project Consortium, 2012) which represents the response variable. The predictor matrix consists of predicted binding sites for each TF, and additional variables, such as G+C composition, can also be included. The results can be visualized as stability paths (Suppl. Fig. 2), that show the selection probability for each motif as a function of the regularization steps, or as the final selection probabilities (Fig. 1C) combined with a sign to indicate if a motif correlates positively or negatively with changes in accessibility.

The illustrating examples and datasets above are included and described in detail in the package vignette. In addition to enrichment- and regression-based motif identification methods, *monaLisa* further provides helpful functions for motif analyses, including functions to predict motif matches and calculate similarity between motifs.

## Summary

*monaLisa* is an R/Bioconductor package for motif analyses applicable to sequences with associated numerical data. Regulatory motifs explaining the observations can be identified using two complementary approaches. *monaLisa* requires no additional software tools and can be easily integrated with other Bioconductor packages for seamless analyses in R.

## Acknowledgements

We would like to thank the members of the Rijli, Schübeler and Stadler groups, Luca Giorgetti, Florian Geier and our colleagues from the Novartis Institutes for Biomedical Research for suggestions on the software. Research in the groups of the authors is supported by the Novartis Research Foundation. DM was supported by the Swiss National Science Foundation (grant 31003A 175776 to FMR). DS furthermore acknowledges support from the Swiss National Science Foundation (310030B 176394) and the European Research Council under the European Union’s (EU) Horizon 2020 research and innovation program grant agreements (ReadMe-667951 and DNAaccess-884664).

## Supplementary Figures

**Supplementary Figure 1:**
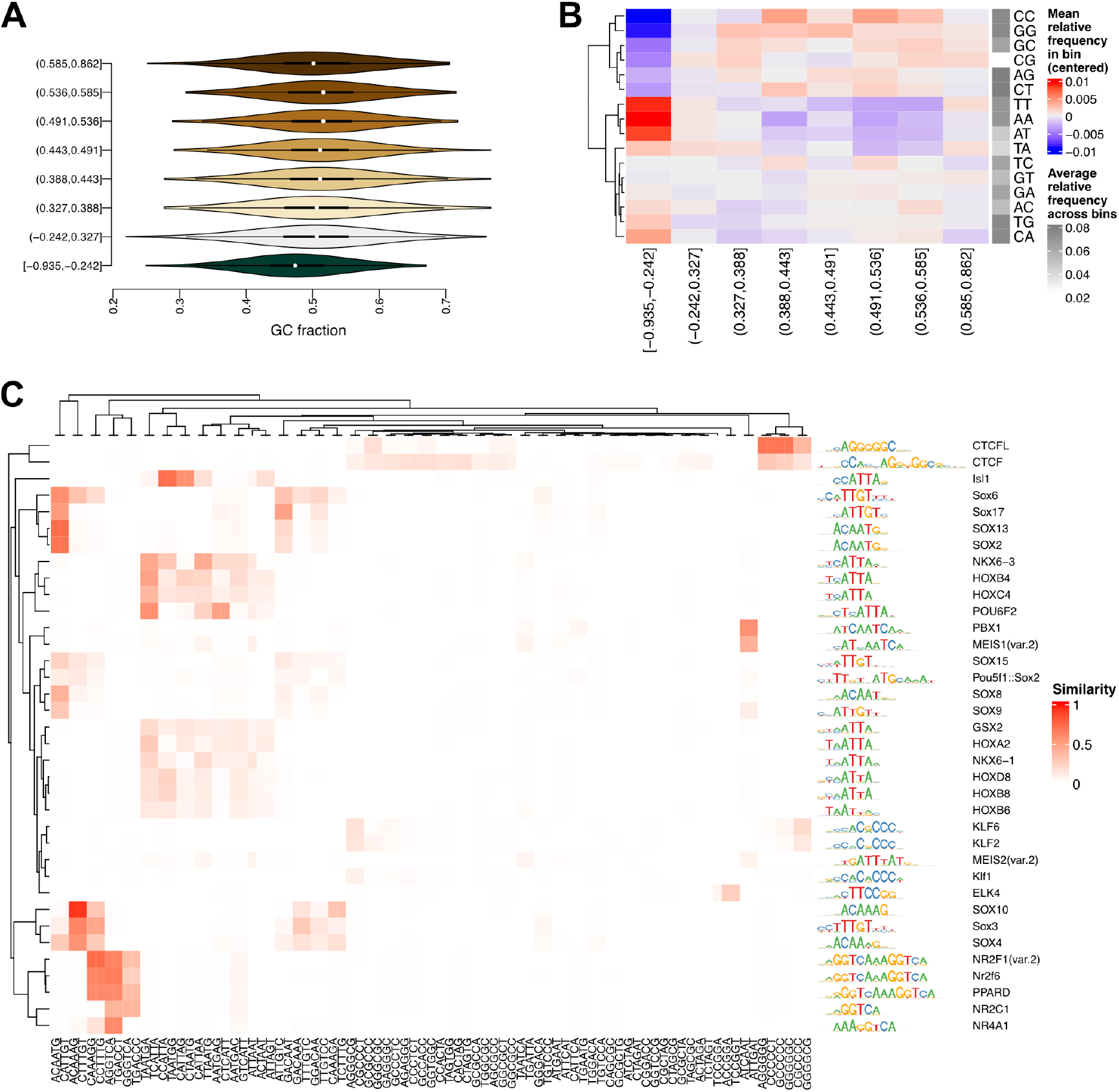
Binned enrichment analysis of methylation changes between embryonic stem cells and neuronal progenitors. **A** Distributions of the fraction of G+C bases for sequences in each bin. **B** Differences of dinucleotide frequencies for sequences in each bin, relative to the mean over all bins. **C** Similarities between enriched motifs (rows) and k-mers (columns). Potential gaps in the motif analysis can be identified as k-mers without similar motifs.

**Supplementary Figure 2:**
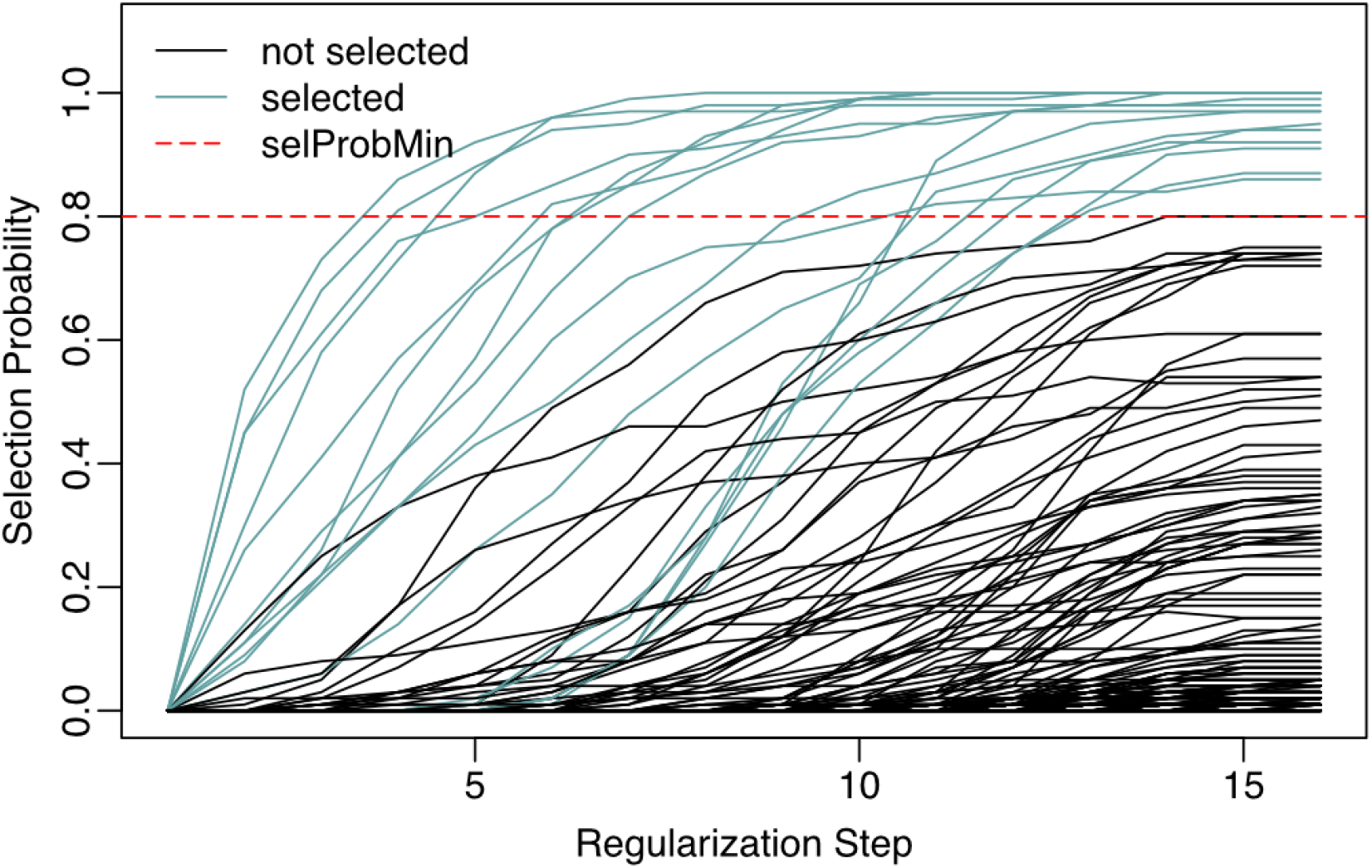
Regression-based analysis of accessibility changes between liver and lung. Stability paths showing the selection probability for each motif as a function of the regularization step, with the legend indicating the colors of selected motifs and the selection probability cutoff.

